# Microbiome ssRNA as an environmental cue to activate TLR13-dependent tissue-protective programs in CD5L^hi^ hepatic macrophages

**DOI:** 10.1101/2023.05.11.540294

**Authors:** Ryota Sato, Kaiwen Liu, Takuma Shibata, Katsuaki Hoshino, Kiyoshi Yamaguchi, Toru Miyazaki, Ryosuke Hiranuma, Ryutaro Fukui, Yuji Motoi, Yuri Fukuda-Ohta, Yun Zhang, Tatjana Reuter, Yuko Ishida, Toshikazu Kondo, Tomoki Chiba, Hiroshi Asahara, Masato Taoka, Yoshio Yamauchi, Toshiaki Isobe, Tsuneyasu Kaisho, Yoichi Furukawa, Eicke Latz, Kensuke Miyake

## Abstract

Hepatic macrophages maintain liver homeostasis, but little is known about the signals that activate the hepatoprotective programs within macrophages. Here, we show that toll-like receptor 13 (TLR13), a sensor of bacterial 23S ribosomal RNA (rRNA), senses microbiome RNAs to drive tissue-protective responses in CD5L^hi^ hepatic macrophages. Splenomegaly and hepatomegaly developed in the absence of the endosomal RNase, RNaseT2, via TLR13-dependent macrophage proliferation. Furthermore, TLR13 in hepatic Ly6C^lo^ macrophages activated the transcription factors LXRα and MafB, leading to expression of tissue-clearance molecules, such as CD5L, C1qb, and Axl. Consequently, *Rnaset2*^−/−^ mice developed resistance to acute liver injury caused by challenges with acetaminophen and lipopolysaccharide + D-galactosamine. TLR13 responses in *Rnaset2*^−/−^ mice were impaired by antibiotics, suggesting that TLR13 were activated by microbiome rRNAs, which was detected in the sera and hepatic macrophages. Repeated administration of wild-type mice with the TLR13 ligand, rather than other TLR ligands, selectively increased the number of Kupffer cells, which expressed immunoregulatory and tissue-clearance genes as hepatic macrophages in *Rnaset2*^−/−^ mice did. Our results suggest that microbiome ssRNA serves as an environmental cue for initiating tissue-protective TLR13 responses in hepatic macrophages.

**Graphical Abstract:** In the absence of an endosomal RNase, RNase T2, microbiome RNAs circulating in the vasculature activate TLR13 in hepatic macrophages to drive hepatoprotective responses through expression of immunoregulatory and tissue-clearance molecules. Consequently, mice lacking RNase T2 are resistant against acute liver injuries caused by acetaminophen and LPS + D-galactosamine.

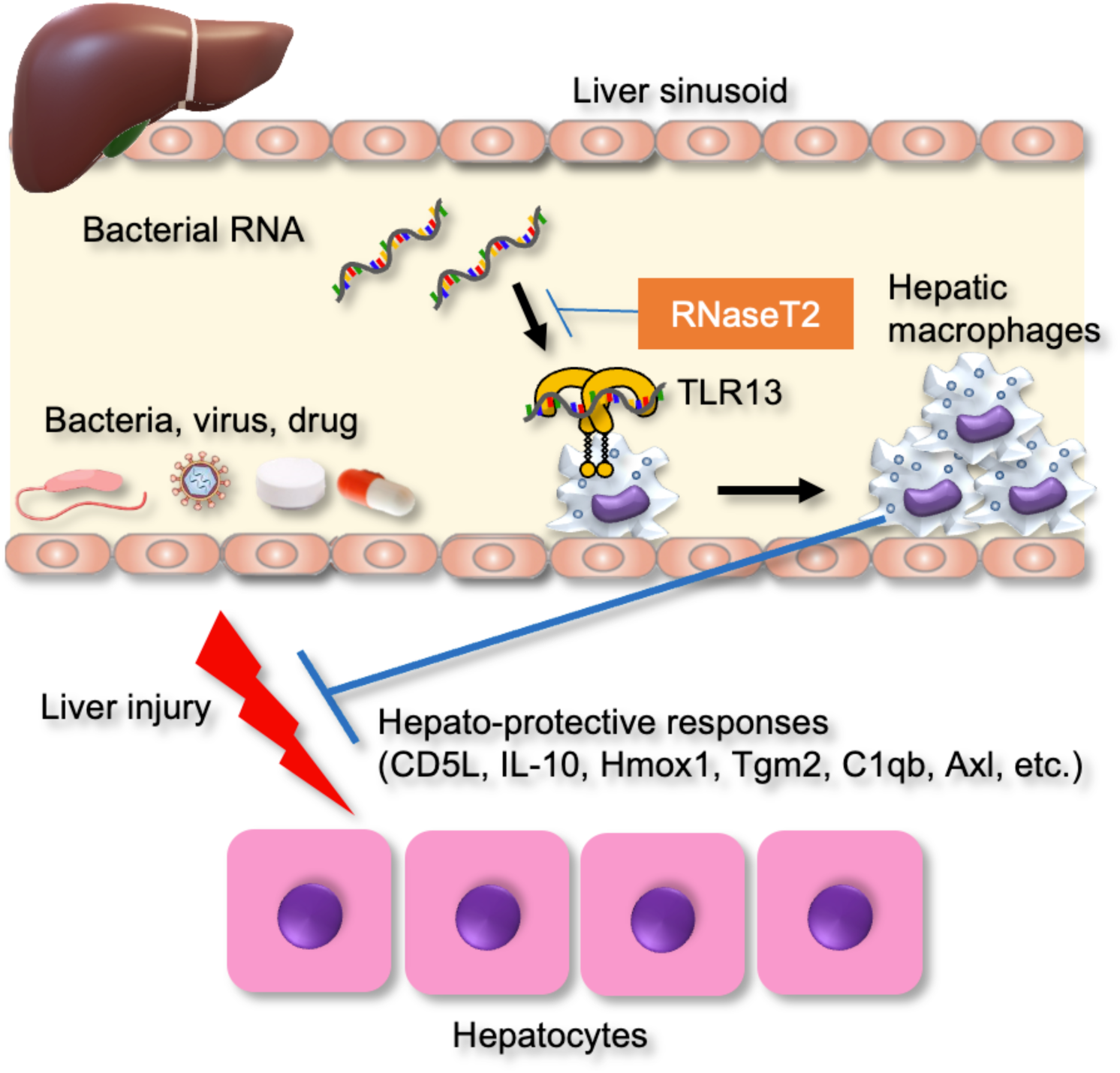

## Introduction

The liver is exposed to blood from the gut, and intestinal bacteria occasionally enter the portal system; hepatic macrophages then clear the bacteria to prevent their dissemination ^1^. In addition to bacteria, drugs are also absorbed from the gut and detoxified by hepatocytes. However, if drug metabolites accumulate owing to excessive drug intake, hepatocytes die and release danger signals ^2, 3^. Hepatic macrophages clear the released danger signals to avoid inflammation ^3^. At least three macrophage subsets protect the liver through clearance. Kupffer cells (KCs) are liver-resident macrophages originating from the yolk sac and replicating in the liver ^4^; they reside in the sinusoidal lumen and clear bacteria, aged neutrophils, and senescent platelets ^5^. Bone marrow-derived Ly6C^hi^ CX3CR1^lo^ monocytes infiltrate the liver upon acetaminophen (APAP) challenge and differentiate into Ly6C^lo^ CX3CR1^hi^ macrophages, promoting clearance and tissue repair ^6^. Peritoneal macrophages also invade the injured liver through the mesothelium and differentiate into alternatively activated macrophages to promote the clearance of dead cells ^7^.

RNAs serve as immune codes that inform immune cells about pathogen invasion and tissue damage ^8, 9^. Extracellular and cytoplasmic RNAs are transported to and degraded in the endosomal compartment, where RNA-sensing toll-like receptors (TLRs), such as TLR3, TLR7, TLR8, and TLR13, reside ^9, 10, 11^. TLR3 responds to double-stranded RNAs (dsRNAs), whereas TLR13 senses single-stranded RNAs (ssRNAs) derived from bacterial 23S ribosomal RNA (rRNA) ^12, 13^. TLR7 and TLR8 sense RNA degradation products that are a combination of nucleosides and oligoribonucleotides (ORNs) ^14–17^. Endosomal RNA degradation affects the response of RNA-sensing TLRs. The endosomal RNase, RNase T2, degrades double-stranded RNA (dsRNA), thereby negatively regulating TLR3 responses ^18^. In contrast, ssRNA degradation by RNase T2 generates ligands for mouse TLR7 and human TLR8 ^18–20^. Splenomegaly, hepatomegaly, and neuroinflammation develop in *Rnaset2*^−/−^ mice ^21^. Little is known, however, about the role of RNA-sensing TLRs in the pathologies of *Rnaset2*^−/−^ mice.

Here, we found that TLR13 drove splenomegaly and hepatomegaly in *Rnaset2*^−/−^ mice. Splenic and hepatic monocytes/macrophages in *Rnaset2*^−/−^ mice TLR13- dependently proliferated and antibiotics inhibited the macrophages proliferation, suggesting that TLR13 was activated by bacterial rRNAs. TLR13 in hepatic macrophages activated tissue-protective responses instead of inflammatory responses through activation of the transcription factors LXRα and MafB, which upregulated the expression of tissue-clearance molecules, such as CD5L, Tgm2, C1qb, and Axl. Consequently, the *Rnaset2*^−/−^ mice were found to be resistant to liver-damaging challenges with APAP and lipopolysaccharide (LPS)+D-galactosamine (D-Gal). In wild-type mice, the TLR13 ligand specifically increased the percentage of KCs, which, like *Rnaset2*^−/−^ hepatic macrophages, expressed tissue-clearance genes. These results suggest that microbiota RNAs serve as an environmental cue for activating tissue-protective TLR13 responses in hepatic macrophages.

## Results

### TLR13-dependent accumulation of monocyte/macrophage in *Rnaset2*^−/−^ mice

Mouse RNase T2 is encoded by two genes, *Rnaset2a* and *Rnaset2b*. To study the role of RNase T2 *in vivo*, previously generated *Rnaset2a*^−/−^ *Rnaset2b*^−/−^ mice were used ^18^; this strain is here described as *Rnaset2*^−/−^ or *Rt2*^−/−^ mice. The mice were born at Mendelian ratios and grew normally. Consistent with a previous report ^21^, splenomegaly developed in *Rnaset2*^−/−^ mice (Fig. 1A), and the proportions of Ly6C^hi^ and Ly6C^lo^ macrophages and red pulp macrophages were increased in their spleens (Fig. 1, B–E). Although the percentages of splenic T and B cells did not increase (Fig. S1A), autoantibodies against RNA-associated antigens (such as Sm and SSA) but not against dsDNA were produced in *Rnaset2*^−/−^ mice (Fig. S1B, S1C). The percentages of splenic conventional dendritic cells (cDCs) and plasmacytoid DCs (pDCs) were not altered (Fig. S1D). In peripheral blood, the counts of both Ly6C^hi^ and Ly6C^lo^ monocytes were increased (Fig. S1E and S1F), whereas those of platelet decreased and macrocytic anemia developed (Fig. 1F; Fig. S1G). Hepatomegaly also developed in *Rnaset2*^−/−^ mice (Fig. 1G). CD11b expression was upregulated in *Rnaset2*^−/−^ hepatic macrophages (Fig. 1H), among which the percentages of Ly6C^lo^ macrophages predominantly increased (Fig. 1I). In addition to the spleen and liver, F4/80^+^ macrophages also infiltrated the brain, lungs, and kidneys (Fig. 1J).

**Fig. 1.**
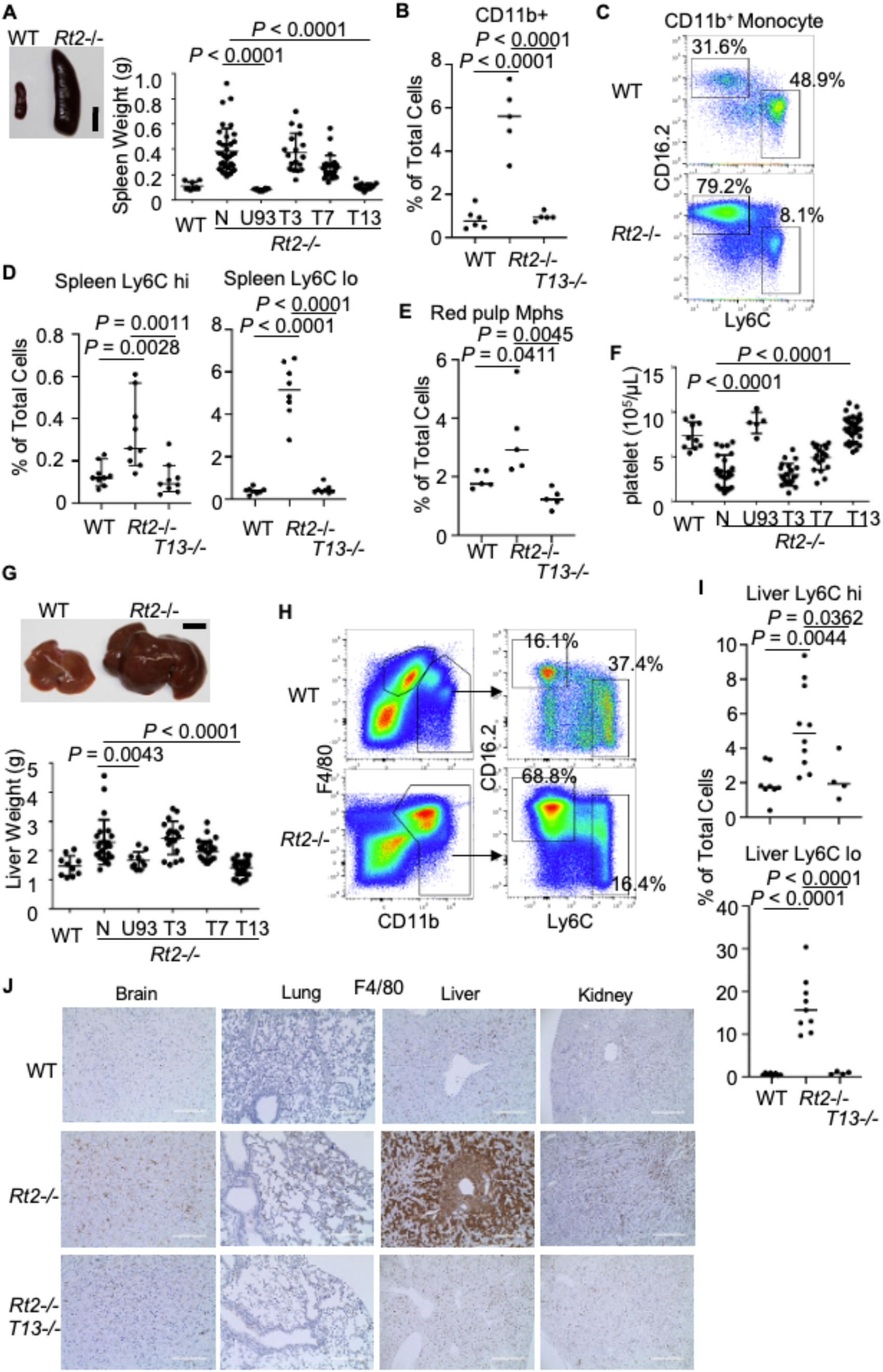
TLR13-dependent macrophage accumulation in *Rnaset2*^−/−^ mice. (A) Spleen photograph and splenic weights of WT (wild-type), *Rt2*^−/−^, *Rt2*^−/−^ *Unc93b1*^−/−^, *Rt2*^−/−^ *Tlr3*^−/−^, *Rt2*^−/−^ *Tlr7*^−/−^, and *Rt2*^−/−^ *Tlr13*^−/−^ mice (n > 8). (B) Percentages of splenic CD11b^+^ macrophages in WT, *Rt2*^−/−^, and *Rt2*^−/−^ *Tlr13*^−/−^ mice (n > 5). (C) Dot plots show the percentages of Ly6C^hi^ CD16.2^lo^ classical monocytes and Ly6C^lo^ CD16.2^hi^ nonclassical monocytes in the spleen of WT and *Rt2*^−/−^ mice. (D) Percentages of splenic Ly6C^hi^ and Ly6C^lo^ monocytes in indicated mice (n > 8). (E) Percentages of F4/80^+^ CD11b^lo^ red pulp macrophages in the spleen of indicated mice (n > 5). (F) Platelet counts of indicated mice (n > 8). (G) Liver photograph and liver weights of indicated mice (n > 10). (H, I) Dot plots show the percentages of Ly6C^hi^ CD16.2^lo^ and Ly6C^lo^ CD16.2^hi^ macrophages in indicated mice (n > 8). (J) Immunohistochemistry showing F4/80 expression in the brain, lung, liver, and kidney of indicated mice. Scale bar, 200 μm.

RNase T2 is localized to endosomes ^18^; therefore, its deficiency causes RNA accumulation in the endosomal compartment ^22, 23^. We hypothesized that monocytosis is caused by the activation of RNA-sensing TLR in *Rnaset2* ^−/−^ mice. Because Unc93b1 is required for all RNA-sensing TLRs ^24^, we first generated *Rnaset2* ^−/−^ *Unc93b1*^−/−^ mice and found that they did not develop splenomegaly, thrombocytopenia, and hepatomegaly (Fig. 1A, 1F, and 1G). We then crossed *Rnaset2*^−/−^ mice with *Tlr3*^−/−^*, Tlr7*^−/−^, or the newly established *Tlr13*^−/−^ mice (Fig. S2). Similar to *Rnaset2*^−/−^ *Unc93b1*^−/−^ mice, splenomegaly, thrombocytopenia, and hepatomegaly were not observed in *Rnaset2*^−/−^ *Tlr13*^−/−^ mice (Fig. 1; Fig. S1). These results suggest that TLR13 activation drove all the phenotypes in *Rnaset2* ^−/−^ mice.

### TLR13-dependent monocyte proliferation in *Rnaset2*^−/−^ mice

Transcriptome analyses of macrophages from *Rnaset2* ^−/−^ and wild-type mice were performed. Compared to the genes expressed in CD11b^+^ macrophages from wild-type mice, 306 and 444 genes were upregulated and downregulated more than 1.5-fold, respectively, in splenic Ly6C^hi^ monocytes from *Rnaset2* ^−/−^ mice (Fig. S3A). Compared to the genes in wild-type CD11b^+^ macrophages, 537 and 217 genes were upregulated and downregulated > 1.5-fold, respectively, in Ly6C^lo^ macrophages from *Rnaset2* ^−/−^ mice (Fig. S3B). Gene set enrichment analysis (GSEA) revealed that proliferation-associated hallmarks, such as “E2F targets”, “G2M checkpoint”, “MYC targets V1”, “mitotic spindle”, and “mTORC1 signaling” were positively enriched in both Ly6C^hi^ and Ly6C^lo^ splenic monocytes (Fig. 2A and 2B), suggesting that these monocytes proliferated in *Rnaset2* ^−/−^ mice. Consistent with this, the expression of the proliferation-associated antigen Ki67 was upregulated in both Ly6C^hi^ and Ly6C^lo^ macrophages from *Rnaset2* ^−/−^ mice (Fig. 2C; Fig. S3C). Uptake of the thymidine analog 5-ethynyl-2’-deoxyuridine (EdU) and DNA content revealed that the percentages of Ly6C^hi^ and Ly6C^lo^ macrophages in the S phase were increased in a TLR13-dependent manner in *Rnaset2* ^−/−^ mice (Fig. 2D to 2G). We also performed transcriptome analyses and proliferation assays of hepatic Ly6C^hi^ and Ly6C^lo^ macrophages (Fig. S3D and S3E, Fig. 2H). Macrophages in the S phase accounted for approximately 0.5–1% of Ly6C^hi^ and Ly6C^lo^ hepatic macrophages in *Rnaset2*^−/−^ mice but less than 0.15% of CD11b^hi^ and CD11b^lo^ hepatic macrophages in wild-type mice (Fid. 2H). These results demonstrate that TLR13 drives macrophage proliferation in the spleen and liver of *Rnaset2* ^−/−^ mice.

**Fig. 2.**
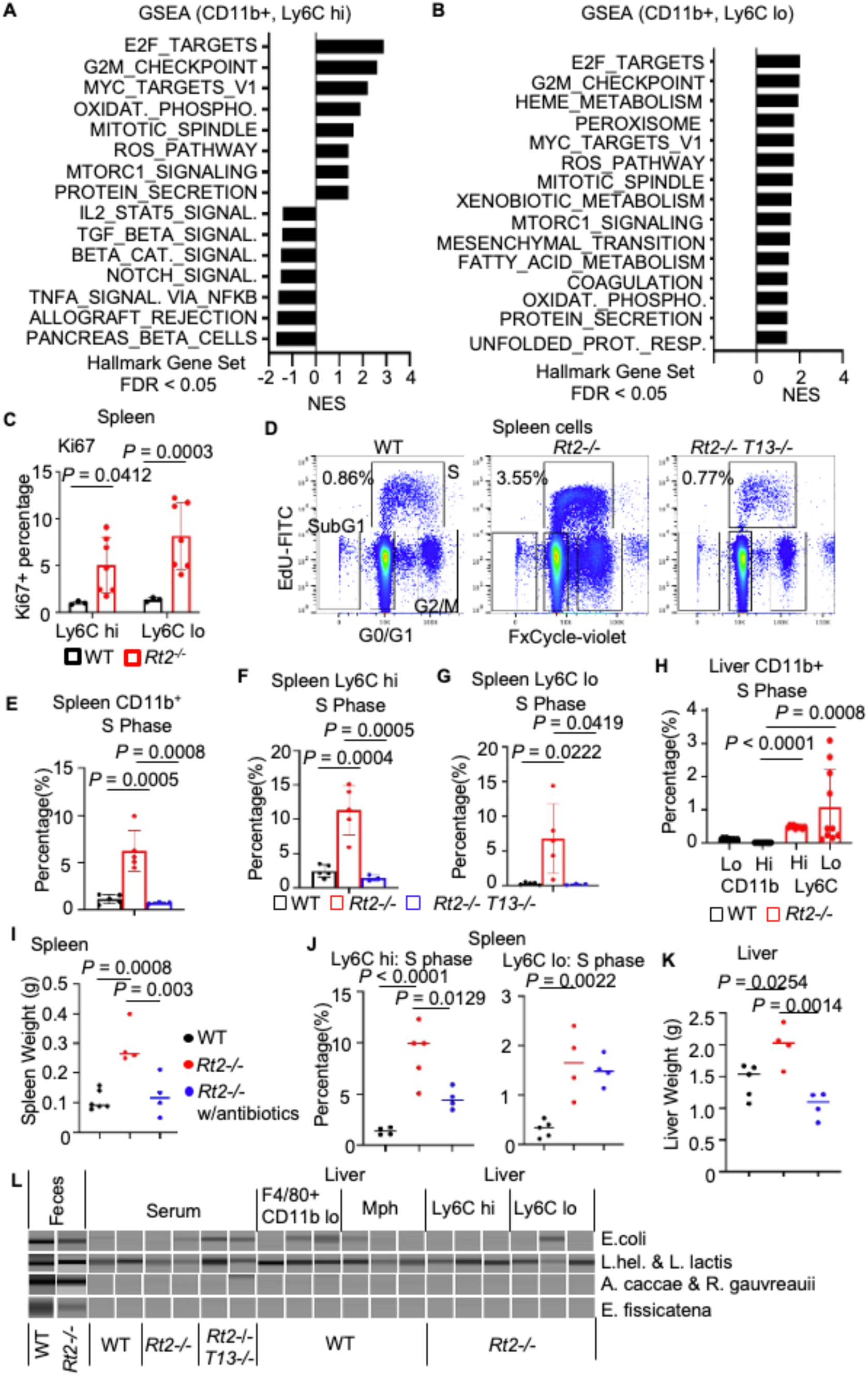
TLR13 drives antibiotic-sensitive monocyte proliferation. (A, B) Transcriptome analyses of splenic Ly6Chi (A) and Ly6Clo (B) macrophages. Gene set enrichment analysis of > 1.5-fold changes in gene expression in Ly6Chi (A) and Ly6Clo (B) splenic macrophages from Rnaset2–/– mice, compared to that in their wild-type counterpart macrophages. The bars indicate normalized enrichment scores (NESs) of hallmarks with FDR < 0.05. (C) Dots show the percentages of Ki67+ cells in Ly6Chi and Ly6Clo splenic macrophages from indicated mice. The results were obtained using FACS (n = 7). (D) Dot plots show EdU uptake and the DNA content (FxCycle violet) in splenic macrophages from indicated mice. The percentages of the S phase are also shown. (E–G) Percentages of cells at S phase in CD11b+ (E), Ly6Chi (F), and Ly6Clo (G) splenic macrophages from indicated mice (n = 5). (H) Percentage of cells at S phase in F4/80+ CD11blo and F4/80+ CD11bhi macrophages from WT mice, and CD11bhi Ly6Chi and CD11bhi Ly6Clo macrophages from Rnaset2–/– mice (n = 10). (I) Dot plots show spleen weights of WT and Rnaset2–/– mice with or without antibiotic treatment from 3 to 6 wk of age. (n > 4). (J) Dot plots show the percentages of S phase in Ly6Chi and Ly6Clo splenic monocytes from indicated mice (n > 4). (K) Dot plots show the liver weights of indicated mice (n > 4). (L) PCR amplification of bacterial 23S rRNA from indicated bacteria. PCR templates were fecal DNA, serum RNAs, and macrophage RNAs. Each panel shows the signal from each mouse.

### TLR13 responses are inhibited by antibiotics

We next focused on TLR13 ligands in *Rnaset2*^−/−^ mice. Although TLR13 was previously found to respond to bacterial 23S rRNA ^12^, hepatomegaly and splenomegaly developed in unperturbed *Rnaset2*^−/−^ mice, suggesting that TLR13 ligands originate from the intestinal microbiota. To study the role of the intestinal microbiome in TLR13-dependent macrophage accumulation, *Rnaset2*^−/−^ mice were treated with a cocktail of antibiotics, namely metronidazole, neomycin, ampicillin, and vancomycin, for 3–6 wk of age. Antibiotics ameliorated splenomegaly by downregulating the proliferation of Ly6C^hi^ macrophages, but not of Ly6C^lo^ macrophages (Fig. 2I and 2J). Hepatomegaly did not develop (Fig. 2K), strongly suggesting that TLR13 in *Rnaset2* ^−/−^ mice was activated by microbiota 23S rRNA. From the database, the consensus sequence in 23S rRNA that activates TLR13 was observed in bacteria, including *Escherichia coli, Lactobacillus helveticus/Lactococcus lactis, Anaerostipes caccae/Ruminococcus gauvreauii,* and *Eubacterium fissicatena* ^25^. PCR detected 23S rRNA sequences from all these bacteria in fecal DNAs from wild-type and *Rnaset2*^−/−^ mice (Fig. 2L), suggesting that all these bacteria reside in the gut. Additionally, 23S rRNAs from *E. coli* and *Lactobacillus helveticus/Lactococcus lactis* were detected in RNAs from serum and hepatic macrophages from wild-type and *Rnaset2* ^−/−^ mice. These results suggest that 23S rRNAs from *E. coli* and *Lactobacillus helveticus/Lactococcus lactis* constitutively translocate into the circulation and act on hepatic macrophages in wild-type and *Rnaset2*^−/−^ mice.

### TLR13 activates the LXRα–CD5L axis in response to the microbiome

We further analyzed the gene expression profiles of splenic and hepatic Ly6C^lo^ macrophages using the search engine “Enrichr” ^26^. The analyses based on the “All RNA-seq and ChIP-seq sample and signature search 4 (ARCHS4)” database suggested activation of transcription factors such as LXR and MafB in hepatic Ly6C^lo^ macrophages from *Rnaset2* ^−/−^ mice (Fig. 3A). The upregulated genes in these macrophages included LXRα target genes, such as *LXRα* itself, *Cd5l*, transglutaminase 2 (*Tgm2*)^27^ (Fig. 3B), and MafB target genes, such as *C1qb* and *Axl* (Fig. 3C) ^27–31^. Fluorescence-activated single cell sorting (FACS) analyses showed that the protein expression levels of LXRα, CD5L, and Axl in hepatic Ly6C^lo^ macrophages from *Rnaset2*^−/−^ mice were increased in a TLR13-dependent manner (Fig. 3D). Although CD5L is a secretory protein, we could detect CD5L protein within macrophages, using membrane-permeabilized staining. CD5L protein levels in circulation and the liver increased in a TLR13-dependent manner (Fig. 3E, 3F). Antibiotics significantly reduced the TLR13-dependent increases of LXRα, CD5L, and Axl in Ly6C^lo^ macrophages (Fig. 3G) and of serum CD5L (Fig. 3H). CD5L protein expression in hepatic Ly6C^lo^ macrophages was downregulated by administration of the LXRα antagonist GSK2033 (Fig. 3I), suggesting that CD5L expression in *Rnaset2* ^−/−^ Ly6C^lo^ hepatic macrophages depends on LXRα activation. In the spleen, we detected considerably smaller increases in the mRNA and protein expression of LXRα and CD5L in *Rnaset2* ^−/−^ Ly6C^lo^ macrophages, but this was not seen for Axl (Fig. S4A, S4B). These differences between the spleen and the liver are consistent with the RNA-seq results that revealed that LXRα and MafB were more strongly activated in hepatic macrophages than in splenic macrophages (Fig. 3A). Together, these results suggest that TLR13 in hepatic Ly6C^lo^ macrophages senses the microbiome RNA to activate the transcription factors LXRα and MafB in *Rnaset2* ^−/−^ mice.

**Fig. 3.**
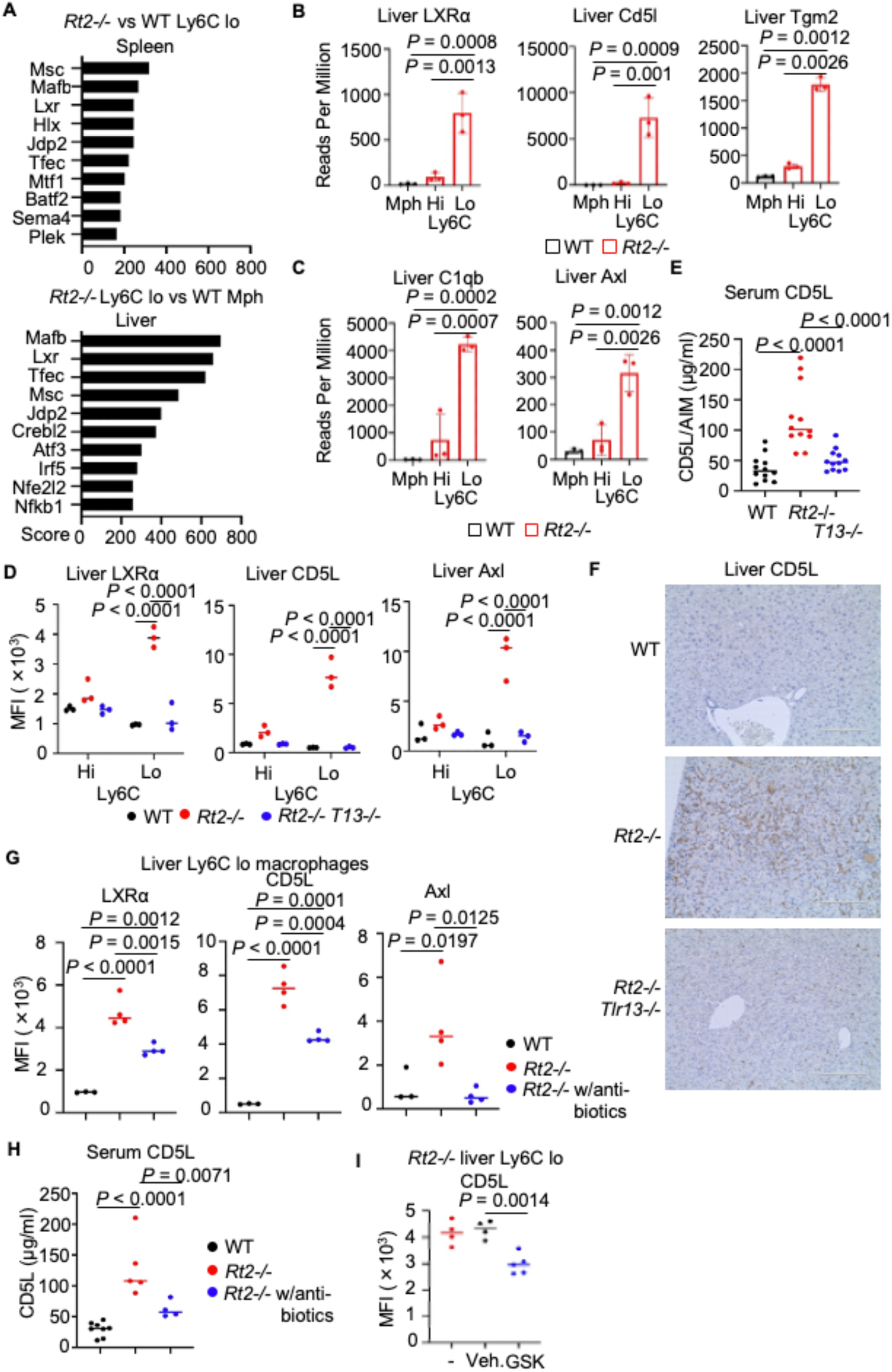
TLR13 activates LXRα and Mafb in *Rnaset2^−/−^* hepatic macrophages. (A) Enrichr analyses to compare *Rnaset2^−/−^*with WT Ly6C^lo^ macrophages from the spleen, and *Rnaset2^−/−^*Ly6C^lo^ macrophages with WT CD11b^+^ macrophages from the liver. (B, C) Bars show reads per million (RPM) of indicated genes in CD11b^hi^ macrophage (Mph) from WT mice, and Ly6C^hi^ and Ly6C^lo^ macrophages from *Rnaset2^−/−^*mice. (D) Mean fluorescence intensity of indicated proteins in Ly6C^hi^ and Ly6C^lo^ macrophages from indicated mice. (E) Serum levels of CD5L in indicated mice (n = 4). (F) Immunohistochemistry of CD5L expression in the liver of indicated mice. Scale bar, 200 µm. (G) Dot plots show the mean fluorescence intensity of indicated proteins in Ly6C^lo^ hepatic macrophages from indicated mice. (H) Serum levels of CD5L in indicated mice (n = 12). (I) Mean fluorescence intensity of CD5L in *Rnaset2^−/−^*mice untreated or treated every other day with vehicle or four times with the LXRα antagonist GSK2033.

### The TLR13–LXRα–CD5L axis in hepatic macrophages is tissue-protective

The genes whose expression was upregulated in hepatic macrophages from *Rnaset2*^−/−^ mice encode immunoregulatory and tissue-clearance molecules. For example, CD5L promotes the clearance of bacteria and damage-associated molecules released during tissue injury ^32^. Other molecules whose expression increased, such as Tgm2, C1qb, and Axl, are known to contribute to cell corpse clearance ^29, 30, 33^. Furthermore, the mRNA expression of genes encoding the anti-inflammatory cytokine IL-10 and the antioxidative enzyme heme oxidase (HO-1) was upregulated in hepatic and splenic Ly6C^lo^ macrophages from *Rnaset2* ^−/−^ mice (Fig. S4C) ^34–36^. Serum IL-10 levels increased in a TLR13-dependent manner (Fig. S4D). In contrast, the mRNA expression of genes encoding proinflammatory cytokines, such as TNFα, IL-6, interferon (IFN)β1, and IL1α, were not upregulated in the spleen and liver (Fig. S4E). Exceptionally, the expression of pro-IL-1β in splenic and hepatic macrophages and IL1α in hepatic macrophages was upregulated in macrophages from *Rnaset2* ^−/−^ mice. These results suggest that tissue-clearance and anti-inflammatory programs, but not inflammatory responses, are constitutively activated in *Rnaset2* ^−/−^ mice.

We hypothesized that *Rnaset2* ^−/−^ mice are resistant to acute liver injuries because molecules whose expression is upregulated, such as IL-10, HO-1, LXRα, and Axl, have protective roles against APAP-induced liver injury ^3, 37, 38^. *Rnaset2*^−/−^ mice were intraperitoneally administered APAP at 750 mg/kg, and ∼70% of wild-type mice did not survive the challenge (Fig. 4A). However, all *Rnaset2*^−/−^ mice survived the APAP challenge, but *Rnaset2*^−/−^ *Tlr13*^−/−^ mice were as sensitive to the APAP challenge as the wild-type mice (Fig. 4A and 4 B). Serum alanine aminotransferase (ALT) and aspartate aminotransferase (AST) levels were not elevated in *Rnaset2*^−/−^ mice (Fig. 4C, 4D), which indicated that hepatocytes escaped APAP-induced death. FACS analyses demonstrated that neutrophils failed to infiltrate the liver in *Rnaset2*^−/−^ mice (Fig. 4E) because production of the neutrophil-attracting chemokine CXCL2 was impaired in the liver (Fig. 4F) ^39^. Furthermore, upon APAP challenge, serum IL-6 was not detected in *Rnaset2*^−/−^ mice (Fig. 4G).

**Fig. 4.**
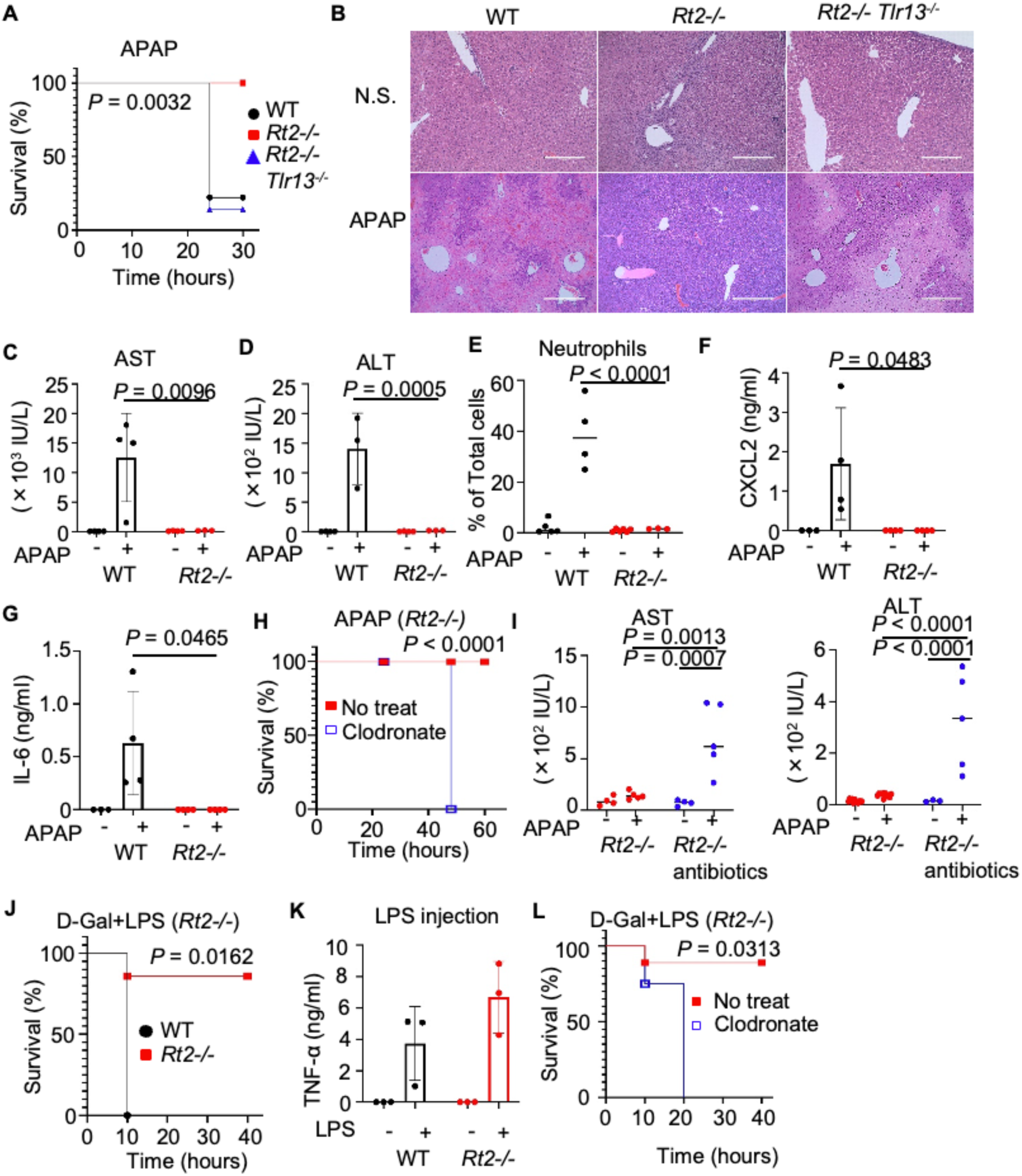
Resistance of *Rnaset2*^−/−^ mice to acute liver injury. (A) Survival curve of indicated mice after APAP challenge at 750 mg/kg (n > 11). (B) HE staining of the liver of indicated mice 24 h after APAP challenge at 500 mg/Kg. Scale bar, 400 µm. (C–G) Dot plots show serum levels of AST (C), ALT (D), CXCL2 (F), and IL-6 (G) and the percentages of neutrophils (E) that infiltrated the liver in indicated mice 24 h after APAP challenge at 500 mg/kg (n = 4). (H) Survival curve of *Rnaset2^−/−^* mice challenged with APAP at 750 mg/kg. Indicated mice had been intravenously administered with clodronate at 25 mg/kg 24 h before APAP challenge (n > 4) (I). Serum levels of AST and ALT in *Rnaset2^−/−^*mice with or without antibiotic treatment (n = 5). (J, L) Survival curve of mice after intraperitoneal administration of LPS and D-Gal at 100 ng/mouse and 12.5 mg/mouse, respectively. Indicated mice intravenously received clodronate at 25 mg/kg 24 h before administration of LPS and D-Gal (L) (n > 5). (K) Serum levels of TNF-α in mice who received intraperitoneal LPS injection at 10 μg/mouse 3 h before blood collection (n = 3).

To examine the role of macrophages in drug resistance, tissue macrophages were depleted with clodronate (Fig. S4F), which made *Rnaset2*^−/−^ mice susceptible to the APAP challenge (Fig. 4H). Antibiotics, which downregulated the expression of LXRα, CD5L, and Axl in Ly6C^lo^ hepatic macrophages (Fig. 3G), also made *Rnaset2*^−/−^ mice sensitive to the APAP challenge, as revealed by the increased levels of serum AST and ALT (Fig. 4I). These results suggest that the liver was made resistant to the APAP challenge by the hepatic macrophages in *Rnaset2*^−/−^ mice.

We next studied the responses to another acute liver injury by administering LPS and D-Gal, which cause TNF-α-dependent hepatocyte apoptosis ^40, 41^. All wild-type mice died within 20 h, whereas >80% of *Rnaset2*^−/−^ mice survived the LPS/D-Gal challenge (Fig. 4J). TNF-α production upon LPS stimulation was not impaired in *Rnaset2*^−/−^ mice (Fig. 4K), but macrophage depletion by clodronate made *Rnaset2*^−/−^ mice susceptible to the LPS/D-Gal challenge (Fig. 4L). These results suggest that TLR13 activation in hepatic macrophages promotes resistance against acute liver injuries.

### TLR13 ligand increases the percentage of KCs in wild-type mice

Splenic and hepatic Ly6C^lo^ macrophages weakly expressed CD5L, whereas peripheral blood Ly6C^lo^ monocytes and hepatic F4/80^+^ CD11b^lo^ macrophages strongly expressed CD5L (Fig. 5A). The subpopulation of hepatic F4/80^+^ CD11b^lo^ macrophages showed stronger CD5L expression and their percentages were increased by the administration of the TLR13 ligand (Fig. 5A), but this was not observed in *Tlr13*^−/−^ mice (Fig. 5B), suggesting that this subpopulation of hepatic macrophages responded to the TLR13 ligand.

**Fig. 5.**
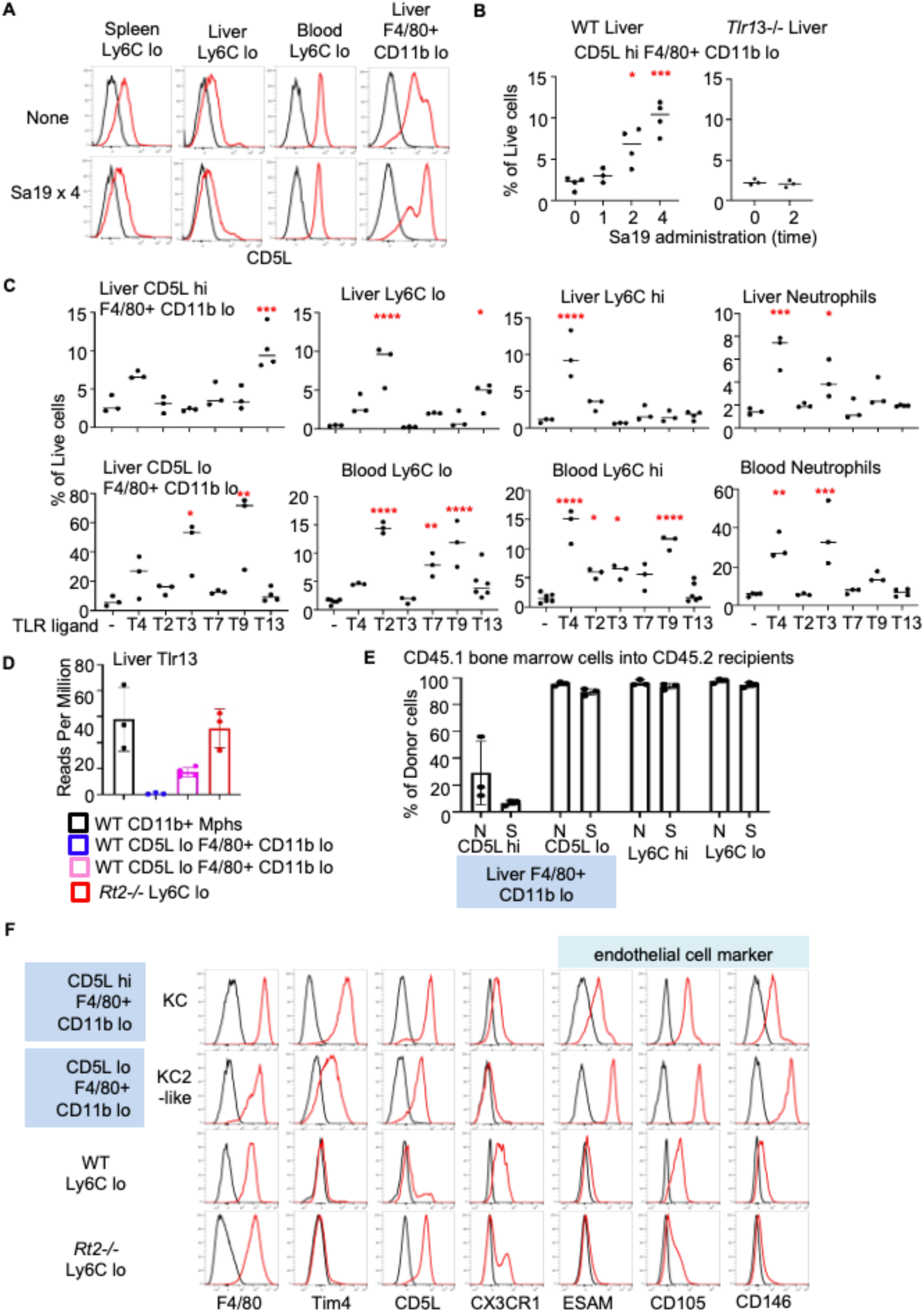
TLR13 ligand increases the percentage of KCs in wild-type mice. (A) Histograms show membrane-permeabilized staining of indicated macrophages with anti-CD5L (red) or isotype-matched (black) antibodies. (B) Dot plots show the percentages of CD5L^hi^ F4/80^+^ CD11b^lo^ hepatic macrophages in WT mice and Tlr13^−/−^ left untreated or treated everyday with the TLR13 ligand Sa19 for 1–4 days (n > 3). (C) WT mice received one of indicated TLR ligands twice. The percentages of indicated macrophages and neutrophils in the liver and blood are shown (n > 3). (D) Dots show reads per million of *TLR13* mRNA in indicated macrophage subsets from indicated mice. (E) Percentages of CD45.1^+^ donor-derived cells in indicated macrophage subsets in bone marrow chimeric wild-type mice that had been irradiated and administered with CD45.1^+^ bone marrow cells. Macrophages were untreated (N) or administered twice with the TLR13 ligand Sa19 (S) (n = 3). (F) Histograms show staining of indicated mice with antibodies against indicated markers (red) or isotype-matched antibody (black).

We compared the TLR13 ligand with other TLR ligands by administering them into wild-type mice. The TLR13 ligand significantly increased the percentages of CD5L^hi^ F4/80^+^ CD11b^lo^ macrophages and weakly increased the percentages of Ly6C^lo^ hepatic macrophages. None of the other TLR ligands increased the percentage of CD5L^hi^ F4/80^+^ CD11b^lo^ macrophages to the same extent as the TLR13 ligand (Fig. 5C; Fig. S5A), but each TLR ligand increased the percentages of other monocyte/macrophage subsets in the liver and blood. For example, LPS strongly increased the percentages of Ly6C^hi^ macrophages and neutrophils in both the liver and blood; TLR2 ligands increased the percentages of hepatic Ly6C^lo^ macrophages and blood Ly6C^lo^ monocytes; TLR3 ligand increased the percentages of CD5L^lo^ F4/80^+^ CD11b^lo^ macrophages, hepatic and blood neutrophils, and blood Ly6C^hi^ monocytes; TLR7 ligand only increased the percentage of blood Ly6C^lo^ monocytes; and TLR9 ligand increased the percentages of CD5L^lo^ F4/80^+^ CD11b^lo^ macrophages and blood monocytes. These differences in the effects of TLR ligands could not be explained by TLR expression (Fig. S5B); for example, *TLR13* mRNA was more highly expressed in hepatic macrophages than in CD5L^hi^ F4/80^+^ CD11b^lo^ macrophages from wild-type mice (Fig. 5D), but the TLR13 ligand more strongly upregulated the percentages of the latter (Fig. 5A). These results suggest that TLR13 responses vary among macrophage subsets.

We focused on the CD5L^hi^ population in F4/80^+^ CD11b^lo^ macrophages. KCs and hepatic macrophages originate from embryos and the bone marrow, respectively ^4^. Bone marrow chimeric mice were used to examine the origin of the CD5L^hi^ subset of F4/80^+^ CD11b^lo^ macrophages. Donor-derived cells replaced more than 90% of the hepatic Ly6C^hi^ and Ly6C^lo^ macrophages and the CD5L^lo^ subset of F4/80^+^ CD11b^lo^ macrophages (Fig. 5E). In contrast, only ∼30% of the CD5L^hi^ subset of F4/80^+^ CD11b^lo^ macrophages was of donor origin (Fig. 5E), and Sa19 administration did not increase the percentage of donor-derived cells among CD5L^hi^ F4/80^+^ CD11b^lo^ macrophages, suggesting that Sa19 did not act on bone marrow-derived hepatic macrophages. CD5L^hi^ F4/80^+^ CD11b^lo^ macrophages were not of BM origin, and highly expressed KC markers, namely F4/80 and Tim4 (Fig. 5F). We hereafter refer to this subset as KCs. The subset of KCs, KC2, characteristically expresses endothelial cell antigens such as ESAM, CD105, and CD146^42^. Antibody analyses showed that CD5L^lo^ F4/80^+^ CD11b^lo^ macrophages expressed these endothelial cell markers as well as KC markers (Fig. 5F and Fig. S6). This bone marrow-derived subset is hereafter referred to as “KC2-like macrophages.”

To further study KCs, KC2-like macrophages, CD11b^+^ macrophages, and *Rnaset2* ^−/−^ Ly6C^lo^ hepatic macrophages, we conducted a transcriptome analysis. Cluster analyses showed that *Rnaset2* ^−/−^ Ly6C^lo^ hepatic macrophages were closer to KCs than to KC2-like or CD11b^+^ hepatic macrophages (Fig. 6A). Consistent with this, tissue-clearance and immunoregulatory genes were more highly expressed in KCs than in KC2-like and CD11b^+^ hepatic macrophages (Fig. 6B). These results suggest that TLR13 drives reparative responses in wild-type KCs.

**Fig. 6.**
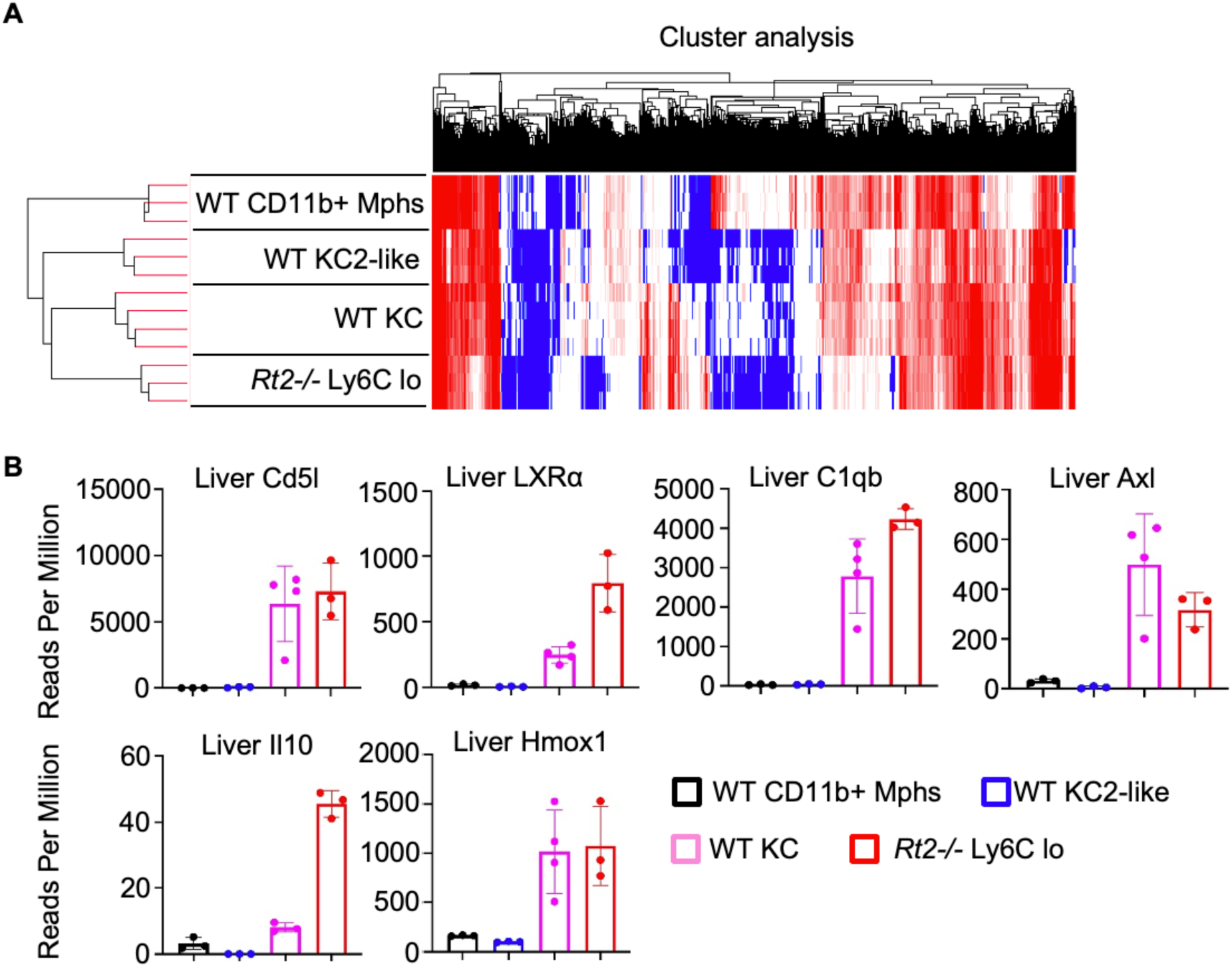
KCs are similar to hepatoprotective *Rnaset2*^−/−^ Ly6C^lo^ macrophages. (A) Cluster analyses of differentially expressed genes between WT CD11b^+^ macrophages and *Rnaset2*^−/−^ Ly6C^lo^ macrophages to compare these macrophage subsets with WT CD5L^hi^ and CD5L^lo^ KCs (n > 3). (B) Bars show reads per million (RPM) of indicated genes in indicated macrophages (n > 3).

## Discussion

The TLR family responds to pathogen components to drive defense responses through the production of proinflammatory cytokines and type I interferons ^11, 43^. In contrast to these well-studied roles, little is known about the role of TLRs in noninflammatory responses, such as proliferation and tissue clearance. In this study, impairing RNA degradation in the endosomal compartment due to RNase T2 deficiency caused lysosomal RNA stress in macrophages, the accumulation of RNAs in endosomes/lysosomes. Lysosomal RNA stress activated TLR13 to drive macrophage proliferation and induce the expression of immunoregulatory and tissue-clearance molecules such as IL-10, HO-1, CD5L, C1qb, and Axl. Consequently, TLR13 responses in macrophages protected *Rnaset2*^−/−^ mice against acute liver injuries. These results suggest that lysosomal RNA stress is a cue to initiate noninflammatory tissue-protective TLR13 responses in macrophages.

Similar TLR13 responses were observed in wild-type mice, where the number of KCs stably expressing tissue-clearance genes, such as *Cd5l, C1qb,* and *Axl*, increased after repeated administration of the TLR13 ligand. KCs expressed other TLRs, such as TLR1, TLR2, TLR3, TLR4/MD-2, TLR5, TLR6, TLR7, and TLR9, but the efficacy of their ligands to increase the number of KCs *in vivo* was much weaker than that of the TLR13 ligand. Such unique noninflammatory TLR13 responses are in part mediated by the transcription factors LXRα and MafB, which induced the transcription of tissue-clearance genes while inhibiting inflammatory responses ^44, 45^.

Hepatic macrophages from *Rnaset2*^−/−^ mice protected hepatocytes from acute liver injury through multiple mechanisms, because many molecules whose expression was upregulated in *Rnaset2*^−/−^ hepatic macrophages, namely IL-10, HO-1, LXRα, and Axl, are reported to protect against the APAP-induced liver injury ^3, 37, 38^. Mechanistically, IL-10 and LXRα directly inhibit immune responses ^34, 44^, whereas CD5L, Tgm2, C1qb, and Axl ameliorate tissue damage by promoting the clearance of damaged cells—the source of danger signals. For example, CD5L plays a protective role against ischemic stroke by promoting the clearance of proinflammatory molecules released from damaged cells, such as peroxiredoxin 1 and HMGB1^46^.

Bacterial 23S rRNAs were found to circulate in the vasculature of unperturbed wild-type mice, but RNA-mediated tissue protection was counterbalanced by endosomal RNA degradation. *Rnaset2*^−/−^ mice revealed that microbiome RNAs positively and negatively regulate TLR responses. In addition to tissue-protective responses and proliferation in macrophages, microbiome RNAs also drove autoantibody production in a TLR13-dependent manner. As B and T cells do not express TLR13, lysosomal RNA stress in DCs is likely to drive autoantibody production against RNA-associated antigens. Despite autoantibody production, glomerulonephritis did not develop in *Rnaset2*^−/−^ mice, which might be explained by impaired TLR7 responses in pDCs and macrophages from *Rnaset2*^−/−^ mice ^18^. Another RNA-sensing TLR, TLR3, is silenced by RNase T2-dependent dsRNA degradation ^18^ and is expected to be hyper-responsive in *Rnaset2*^−/−^ mice. Although TLR3 interacts with dsRNAs from lactic acid bacteria^47^, TLR3 deficiency did not alter monocytosis in *Rnaset2*^−/−^ mice. As the TLR3 ligand poly(I:C) increased the number of blood Ly6C^hi^ monocytes, their increases in *Rnaset2*^−/−^ mice may have been redundantly driven by TLR3.

TLR13 ligation by RNase T2 deficiency activated the transcription factors LXRα and MafB in ly6C^lo^ hepatic macrophages, whereas TLR13 ligation in BM-derived macrophages induces production of proinflammatory cytokines^12^, suggesting that tissue-protective TLR13 responses are restricted to specific macrophage subsets. In wild-type mice, KCs highly expressed CD5L and increased in number upon stimulation with the TLR13 ligand. TLR13 ligation in KCs is likely to activate noninflammatory tissue-protective responses through the LXRα–CD5L axis. KCs increased in number only after the second administration of the TLR13 ligand. Considering that repeated TLR responses induce tolerance in inflammatory responses^48^, repeated TLR13 ligation may influence the downstream signaling pathways to activate LXRα and MafB. Microbiome ssRNAs, which were steadily detected in the vasculature and hepatic macrophages, are likely to continuously act on TLR13 in KCs to activate hepatoprotective programs. High CD5L expression was also seen in peripheral blood Ly6C^lo^ patrolling monocytes, which circulate to clear damaged endothelial cells ^49^. Because the TLR13 ligand failed to increase patrolling monocytes in the vasculature, another environmental cue and another TLR might control clearance programs in this macrophage subset. Mouse KCs are divided into two subsets: KC1 and KC2 ^50^, where the latter’s percentage increases under a high-fat diet and promote diet-induced obesity^50^. Lipids from the gut might be another environmental cue sensed by specific macrophage subsets in the liver and spleen.

In conclusion, we here show that TLR13 in hepatic macrophages senses microbiome ssRNA to activate hepatoprotective responses and that RNase T2 negatively regulates the tissue-protective TLR responses through RNA degradation. Microbiome ssRNA serves as an environmental cue to initiate hepato-protective programs. These results help us understand mechanisms by which homeostasis is dynamically maintained by macrophage responses to environmental cues.

## Methods

### Mice

C57BL/6 mice (sex; male and female, weight; 14-20g) were purchased from Japan SLC, Inc. (Shizuoka, Japan). *Rnaset2a^−/−^ Rnaset2b^−/−^* and *Unc93b1^−/−^*mice were previously described ^18, 51^. In this manuscript, we refer to *Rnaset2a^−/−^ Rnaset2b^−/−^*mice as *Rnaset2^−/−^*mice for simplicity. C57BL/6 *Tlr3^−/−^*and *Tlr7^−/−^* mice were kindly provided by Professor Shizuo Akira (Osaka University, Japan) and have been previously described ^52, 53^. All the animals were housed in SPF facilities at the Institute of Medical Science, University of Tokyo (IMSUT). All animal experiments were approved by the Institutional Animal Care and Use Committee of the IMSUT (#PA17-84, #PA22-43).

### Generation of *Tlr13^−/−^* mice

CMTI-2 (Bruce4) Embryonic Stem (ES) cells were transfected with the vectors targeting the *Tlr13* locus (Fig. S2A), and clones resistant to G418 and ganciclovir were screened for homologous recombination using PCR and confirmed using Southern blot analysis. Targeted ES clones were injected into BALB/c-derived blastocysts to generate chimeric mice, which were mated to obtain *Tlr13*^−/−^ mice. *Tlr13*^−/−^ mice were typed by PCR using primers (Primer#1: 5’-TCGGAAACCTACCCAAGTTAGAGACAC-3’, Primer#2: 5’-TAACTCCTGCAAACTACCCAATCCTTG-3’, Primer#3: 5’-ATCGCCTTCTATCGCCTTCTTGACGAG-3’)

### Reagents

The LXR agonist T0901317 and LXR antagonist GSK2033 were purchased from Selleck Chemicals (Houston, TX, USA), clodronate from Funakoshi (Tokyo, Japan), and acetaminophen (APAP) from TCI Chemicals (Tokyo, Japan). The EdU used in the in vitro proliferation assay was purchased from Tokyo Chemical Industry Co. (Tokyo, Japan). Lipid A purified from *Salmonella minnesota* (Re-595) and lipopolysaccharide (LPS) from *Escherichia coli* (O55:B5) were purchased from Sigma-Aldrich (Merck, Darmstadt, Germany). Pam3CSK4, poly(I:C), and R848 were purchased from InvivoGen (Hong Kong, China). Sa19 (19mer, GsGsAsCsGsGsAsAsAsGsAsCsCsCsCsGsUsGsG) and CpGB ODN1668 (dTsdCsdCsdAsdTsdGsdAsdCsdGsdTsdTsdCsdCsTdsdGsdAsdTsdGsdCsdT), in which ‘s’ depicts a phosphorothioate linkage, were synthesized by FASMAC (Kanagawa, Japan). Recombinant CD5L preparation has been described previously ^54^. Lipofectamine 2000 was purchased from Invitrogen (Thermo Fisher Scientific, Waltham, MA, USA), and DOTAP from Sigma-Aldrich.

### Antibodies

Rat anti-mouse TLR1 monoclonal antibody (mAb) (TR23), rat anti-mouse TLR2 mAb (CB225), mouse anti-mouse TLR5 mAb (ACT5), mouse anti-mouse TLR6 mAb (C1N2), mouse anti-mouse TLR3 mAb (PaT3), mouse anti-mouse TLR7 mAb (A94B10), and mouse anti-mouse TLR9 mAb (J15A7) were established in our laboratory.

Phycoerythrin (PE)-conjugated mouse anti-TLR3 mAb (PaT3), PE-conjugated mouse anti-TLR7 mAb (A94B10), and PE-conjugated mouse anti-TLR9 mAb (J15A7) were purchased from BD Biosciences (Franklin Lakes, NJ, USA). Biotinylated mAbs were prepared using Biotin-XX (Thermo Fisher Scientific). Biotinylated mouse anti-mouse TLR4 mAb (UT49) was provided by Dr. Hiroki Tsukamoto (Fukuoka, Japan). PE rat IgG2a-isotype control antibody and PE mouse IgG2b-d κ isotype control antibody were purchased from BioLegend (Sandiego, CA, USA) and PE mouse IgG1-κ Isotype control antibody from BD Biosciences. Monoclonal anti-mouse CD11b (clone M1/70), CX3CR1 (clone SA011F11), F4/80 (clone BM8), NK1.1 (clone PK136), CD16.2 (clone 9E9), CD3ε (clone 145-2c11), CD19 (clone 6D5), CD11c (clone N418), CD317 (clone 927), CD45.2 (clone 104), CD8 (clone 53-6.7), Ki67 (clone 16A8), Tim4 (clone F31-5G3), ESAM (clone 1G8/ESAM), CD105 (clone MJ7/18), CD59a (clone mCD59.3), CD31 (clone 390), CD146 (clone ME-9F1), CD48 (clone HM48-1), CD11a (clone M17/4), CD24 (clone M1/69), and CD64 (clone X54-5/7.1) antibodies were purchased from BioLegend.

Monoclonal anti-mouse CD49b (clone Hmα2), IA/IE (clone M5/114.15.2), Ly6C (clone HK1.4), and Ly6G (clone 1A8) antibodies were purchased from BD biosciences. Monoclonal anti-mouse CD4 (clone RM4-5) antibody was purchased from Invitrogen and monoclonal anti-mouse Axl antibody (clone 175128) from R&D Systems (Minneapolis, MN, USA). Polyclonal anti-mouse LXRα antibody (#ab3585) was purchased from Abcam (Cambridge, UK). The rabbit anti-mouse CD5L antibody (clone rab1) was provided by Dr. Miyazaki (Tokyo, Japan). The LEGENDScreen Mouse PE Kit was purchased from BioLegend.

### Cell preparation

Blood cells were obtained from mice, using a microtube with EDTA (Erma Inc., Tokyo, Japan). Livers were minced and processed using a gentle MACS Octo Dissociator with Heaters (Miltenyi Biotec, Bergisch Gladbach, Germany). Supernatants were filtered using MACS SmartStrainer (pore size: 100 µM; Miltenyi Biotec) and centrifuged at 300 ×*g* for 10 min. The pellet was resuspended in Debris Removal Solution (Miltenyi Biotec) and centrifuged at 3000 ×*g* for 10 min. The cell pellet was resuspended in RBC lysis buffer (BioLegend).

### Flow cytometry

Cell surface staining for flow cytometric analysis was performed using fluorescence-activated cell sorting (FACS) staining buffer (1 × phosphate-buffered saline [PBS] with 2.5% fetal bovine serum and 0.1% NaN_3_). The prepared cell samples were incubated for 10 min with an unconjugated anti-mouse CD16/32 blocking mAb (clone 95) to prevent nonspecific staining in the staining buffer. The cell samples were then stained with fluorescein-conjugated monoclonal antibodies for 20 min on ice. Stained cells were fixed with BD Cytofix Fixation Buffer (BD Biosciences) for 20 min at 4°C and washed with the staining buffer. For intracellular staining of TLR3, 7, and 9, fixed cells were permeabilized using BD Perm/Wash buffer (BD Biosciences) and incubated with anti-TLR antibody or isotype control IgG1 for 30 min at 4°C. For intracellular staining of CD5L, cell-surface stained cells were fixed and permeabilized using True-Nuclear Transcription Factor Buffer Set (BioLegend), incubated with anti-CD5L antibody for 30 min at 4°C, and washed with True-Nuclear Perm Buffer. The stained cells were incubated with PE-conjugated anti-rabbit IgG for 30 min at 4°C and washed with True-Nuclear Perm Buffer. The stained cells were analyzed using an ID7000 spectral cell analyzer (Sony Biotechnology, San Jose, CA, USA). All data were analyzed using FlowJo software (BD Biosciences).

### Cell sorting

Cell sorting was conducted using the FACS ARIA III Cell Sorter (BD Biosciences). To purify Ly6C^lo^ and Ly6C^high^ splenic monocytes, splenocytes from wild-type and Rnaset2^−/−^ mice were incubated with biotinylated anti-mouse CD3 (clone 145-2C11)/CD19 (clone 6D5)/NK1.1 (clone PK136)/Ly6G (clone aA8)/TER-119/erythroid cells (clone Ter-119), followed by incubation with Streptavidin MicroBeads (Miltenyi Biotec). The magnetically labeled cells were removed using autoMACS (Miltenyi Biotec), and the enriched cells were stained with anti-mouse CD45.2, F4/80, CD11b, Ly6C, CD16.2, NK1.1, and Ly6G mAbs. Ly6C^lo^ CD16.2^high^ and Ly6C^high^ CD16.2^lo^ CD11b^+^NK1.1^−^Ly6G^−^ cell populations were sorted. For sorting of KCs and hepatic macrophages from wild-type mice and Ly6C^lo^ and Ly6C^high^ hepatic macrophages from Rnaset2^−/−^ mice, liver cells were stained with antibodies against CD45.2, F4/80, CD317, CD11b, Ly6C, and CD16.2. F4/80^+^ CD11b^lo^, F4/80^+^ CD11b^hi^, F4/80^+^ CD11b^lo^ CD317^hi^ CD16.2^lo^, and F4/80^+^ CD11b^lo^ CD317^lo^ CD16.2^hi^ cells were sorted as KCs, hepatic macrophages, CD5L^lo^ KCS, and CD5L^high^ KCs, respectively. F4/80^+^ CD11b^hi^ Ly6C^lo^ CD16.2^hi^ and F4/80^+^ CD11b^hi^ Ly6C^lo^ CD16.2^hi^ cells in *Rnaset2*^−/−^ mice were sorted as Ly6C^lo^ hepatic macrophages and Ly6C^high^ hepatic macrophages.

### RNAseq analysis

Total RNA was extracted from sorted cells using RNeasy Mini Kits (Qiagen, Hilden, Germany), and the quality of the RNA was evaluated using an Agilent Bioanalyzer (Agilent Technologies, Santa Clara, CA, USA). Samples with an RNA integrity number value > 7.0 were subjected to library preparation. RNA-seq libraries were prepared with 1 ng of total RNA using the Ion AmpliSeq Transcriptome Mouse Gene Expression kit (Thermo Fisher Scientific) according to the manufacturer’s instructions. The libraries were sequenced with 100-bp single-end reads to a depth of at least 10 million reads per sample on the Ion Proton platform, using an Ion PI Hi-Q Sequencing 200 kit and Ion PI Chip v3 (Thermo Fisher Scientific). The FASTQ files were generated using AmpliSeqRNA plug-in v5.2.0.3 in the Torrent Suite software (v5.2.2; Thermo Fisher Scientific) and analyzed using the TCC-GUI software. Individual sample reads were normalized to relative log expression using the DESeq2 R library. DESeq2 was used to determine the fold changes and *p*-values. Genes showing > 1.5-fold change in expression (adjust *p* < 0.05) were considered significantly altered. To interpret the gene expression profiles, gene set enrichment analysis (GSEA) was performed using GSEA 4.1.0, with the MSigDB hallmark gene sets. Enriched pathways were determined by FDR-adjusted *p*-values < 0.1 To identify the activation transcription factors, over representation analysis was conducted using Enrich R (https://maayanlab.cloud/Enrichr/) with ARCHS4 TFs Coexp.

### Proliferation assay with EdU labelling

*In vitro* proliferation assays were conducted using the Click-iT Plus EdU Alexa Fluor 488 Flow Cytometry Assay Kit (Invitrogen), according to the manufacturer’s instructions. Spleen, liver, and blood samples were collected from the mice. Erythrocytes were then completely lysed using BD Pharm Lyse lysing buffer (BD Biosciences) to collect splenocytes and peripheral blood mononuclear cells (PBMCs). Collected cells were incubated with 1 µg/ml EdU for 1 h. After blocking splenocytes and PBMCs with an anti-CD16/32 (clone 95) mAb, the samples were stained with fluorescent dye-conjugated mAbs. The stained samples were subsequently fixed with BD Cytofix (BD Biosciences) and permeabilized using 1× Click-iT saponin-based permeabilization and washing reagents. Finally, EdU incorporated into the genomic DNA was stained using the Click-iT EdU reaction cocktail. EdU-positive cells were detected using the abovementioned spectral flow cytometer ID7000 (Sony Biotechnology).

### Histological analysis

Mouse tissues were fixed in 20% formalin neutral buffer solution. Fixed kidneys were embedded in paraffin wax for sectioning. Sections were subjected to hematoxylin and eosin (HE) staining or immunohistochemistry for F4/80 and CD5L and visualized using an EVOS microscope (Thermo Fisher Scientific).

### Biochemical test

Sera were collected from mice aged 30–45 weeks. Aspartate amino transferase (AST) and alanine aminotransferase (ALT) levels were measured using the Biochemical automatic analyzer JCA-BM6050 (JEOL Ltd., Tokyo, Japan) in ORIENTAL YEAST Co., Ltd. (Tokyo, Japan)

### Platelet and cell counts

Platelet numbers in PBMCs were analyzed using an automatic hematology analyzer Celltac α (Nihon Kohden, Tokyo, Japan), and cells were counted using an automated cell counter CellDrop BF (DeNovix, Wilmington, DE, USA).

### TLR ligand injection

Each ligand was diluted in PBS. Mice were intravenously administered with lipid A (10 µg/mice), Pam3csk (10 µg/mice), R848 (10 µg/mice), poly(I:C) (100 µg/mice), CpG-B (10 µg/mice with DOTAP), and Sa19 (2 µg/mice with Lipofectoamine) for two consecutive days.

### Establishment of APAP-induced mouse liver injury

Mice were allowed free access to water but not food for 16 h before the APAP challenge. APAP was dissolved in PBS containing 10% DMSO. In preliminary experiments, mice were intraperitoneally administered with an APAP solution at a dose of 250–750 mg/kg. To determine the survival rate, a dose of 750 mg/kg was administered. To evaluate liver injury based on serum AST and ALT levels and neutrophil infiltration into the liver, we selected 500 mg/kg as the APAP dose. For experimental intervention, mice were intravenously administered with rCD5L (400 µg/mice) at the same time as the APAP challenge; clodronate (25 mg/mice) at 16 h before the APAP challenge; or Ly6C^lo^ hepatic macrophages from *Rnaset2*^−/−^ mice (1×10^6^ cells/mice) 16 h before the APAP challenge.

### ELISA

Anti-Sm and anti-SSA/Ro60 antibodies were quantified using an ELISA kit (Alpha Diagnostic International Inc.). Serum levels of anti-double-stranded DNA antibodies were measured using a commercial ELISA kit (FUJIFILM Wako Pure Chemical Corporation, Osaka, Japan), serum IL-10 and CXCL2 levels were measured using a DuoSet ELISA kit (R&D Systems), and serum IL-6 and TNF-α levels were measured using a commercial ELISA kit (Thermo Fisher). Serum CD5L levels were measured using ELISA, as described previously ^54^.

### Data and materials availability

All data, code, and materials used in the analysis must be available in some form to any researcher for purposes of reproducing or extending the analysis. Include a note explaining any restrictions on materials, such as materials transfer agreements (MTAs). Note accession numbers to any data relating to the paper and deposited in a public database; include a brief description of the data set or model with the number. If all data are in the paper and supplementary materials, include the sentence “All data are available in the main text or the supplementary materials.”

## Acknowledgments

We acknowledge the FACS Core laboratory at the Institute of Medical Science in the University of Tokyo for assistance with the cell sorting by FACSAria flow cytometer. We would like to thank Editage (www.editage.com) for English language editing. This work was supported in part by: JSPS/MEXT KAKENHI Grants: JP 21H04800, JP 22H05184, JP 22K19424, and JP 22H05182 to K. M; JP 19K16685 and JP 21K15464 to R. S; JP 19H03451, JP 16K08827 to T.S; JP 17K19568, JP 17H04088, JP 19H04813, JP 20H03505, JP 21K19384, JP 22H05182, JP 22H05187 to T. K; JP 21K08458 to K. H; JP 16K19585 and JP 18K16096 to Y. F-O; and JST CREST (JPMJCR21E4); the Japan Agency for Medical Research and Development (AMED) Grant Number JP 20ek0109385 to T.S; Takeda Science Foundation to R.S and T. K; Daiichi Sankyo Foundation of Life Science to R. S; Mochida Memorial Foundation for Medical and Pharmaceutical Research to R.S and T.S; the Uehara foundation to T. K; Joint Research Project of the Institute of Medical Science at the University of Tokyo; JSPS KAKENHI Grant Number JP 16H06276 (AdAMS); University of TOkyo Pandemic preparedness, Infection and Advanced research Center (UTOPIA); and AMED-LEAP (JP22gm0010006h) to TM.

## Author contributions

RS and KM conceived of and designed the experiments. TS, KH, Y F-O, MO and TK generated mutant mice. MT, YM, and TI conducted biochemical analysis. RS, KY and FY performed the transcriptome analyses. RS performed the *in vivo* and *in vitro* analyses with the help of KL, TS, RH, RF, YM, YZ and TR. RS performed all other experiments in this study and analysed the data for all figures. KH, TM, YI, TK, TC, HA, TK and EL discussed obtained results. RS and KM wrote the paper. All authors have reviewed the manuscript

---

**Fig. S1.**
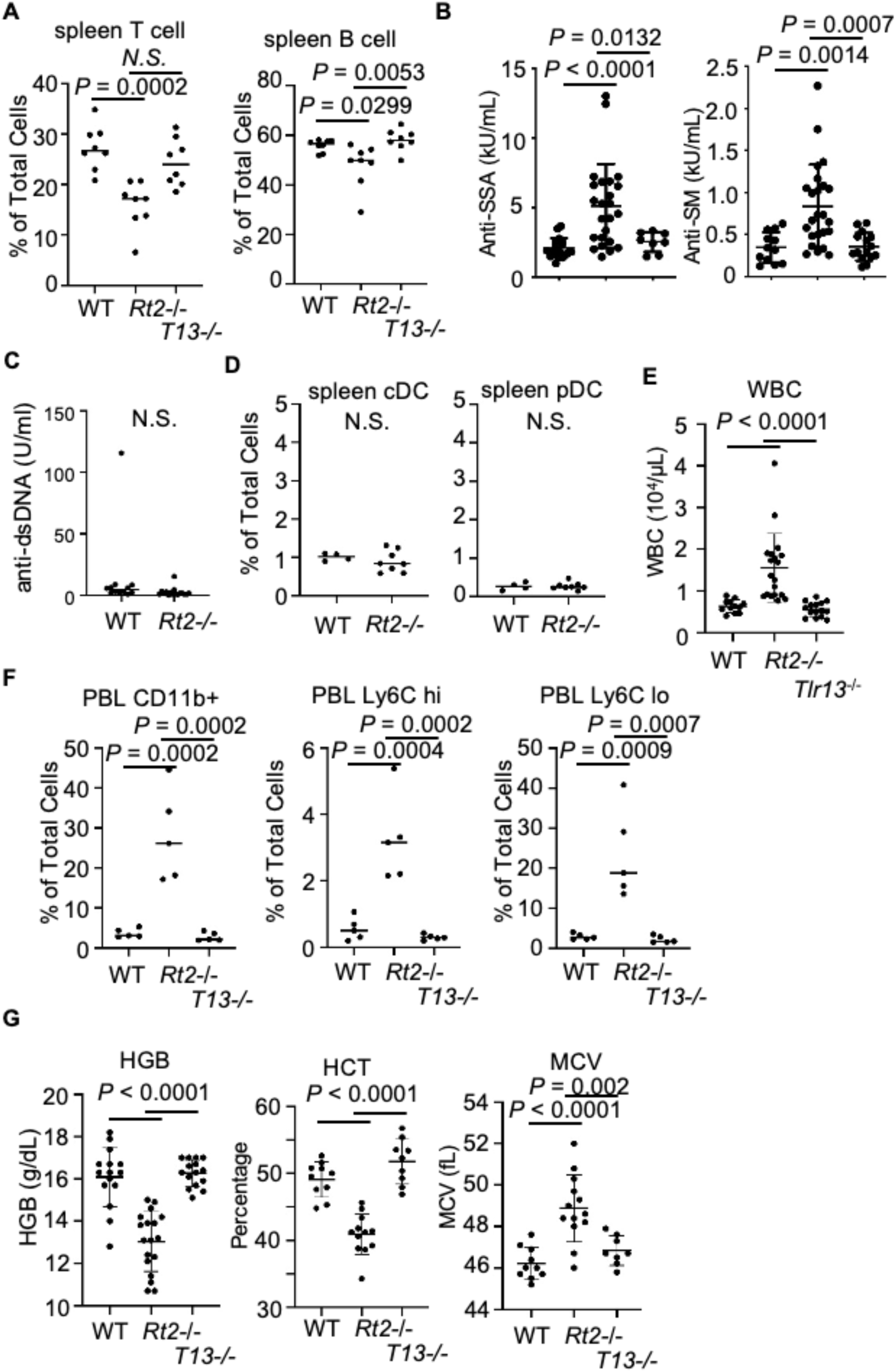
Phenotypes of *Rnaset2^−/−^* mice. (A) Percentages of splenic T cells and B cells in indicated mice (n = 8). (B, C) Serum titres of autoantibodies to SSA, Sm, and dsDNAs in indicated mice (n > 8). (D) Percentages of CD11c^+^ I-A^+^ cDCs and CD11c^+^ PDCA-1^+^ pDCs in the spleen of indicated mice (n > 4). (E, F) Percentages of whole blood cells (E) and Ly6C^hi^ and Ly6C^lo^ monocytes (F) in the blood of indicated mice (n > 5). (G) Hb concentration, hematocrit, and mean corpuscular volume of indicated mice (n > 8).

**Fig. S2.**
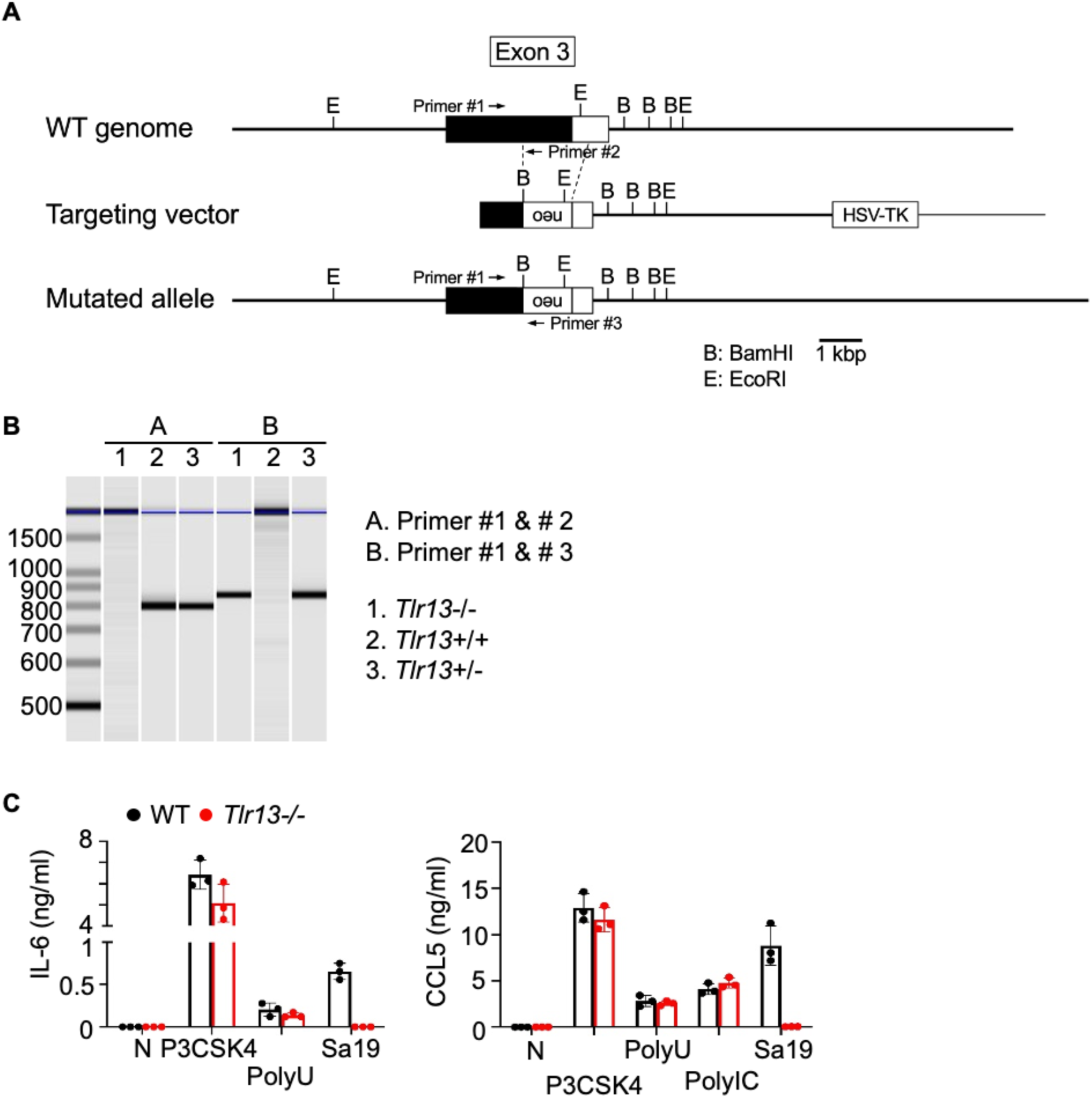
Generation of *Tlr13*^−/−^ mice. (A) Schematic representation of gene targeting. The filled and open boxes represent coding and 3’-untranslated regions of the *Tlr13* gene, respectively. Neo indicates the neomycin resistance gene: B, BamH I; and E, EcoR I. (B) PCR analyses with primers indicated in (A) to determine genotypes of indicated mice. (C) Production of IL-6 and CCL5 by bone marrow-derived macrophages unstimulated or stimulated with indicated TLR ligands. Results represent mean values with SD from triplicates.

**Fig. S3.**
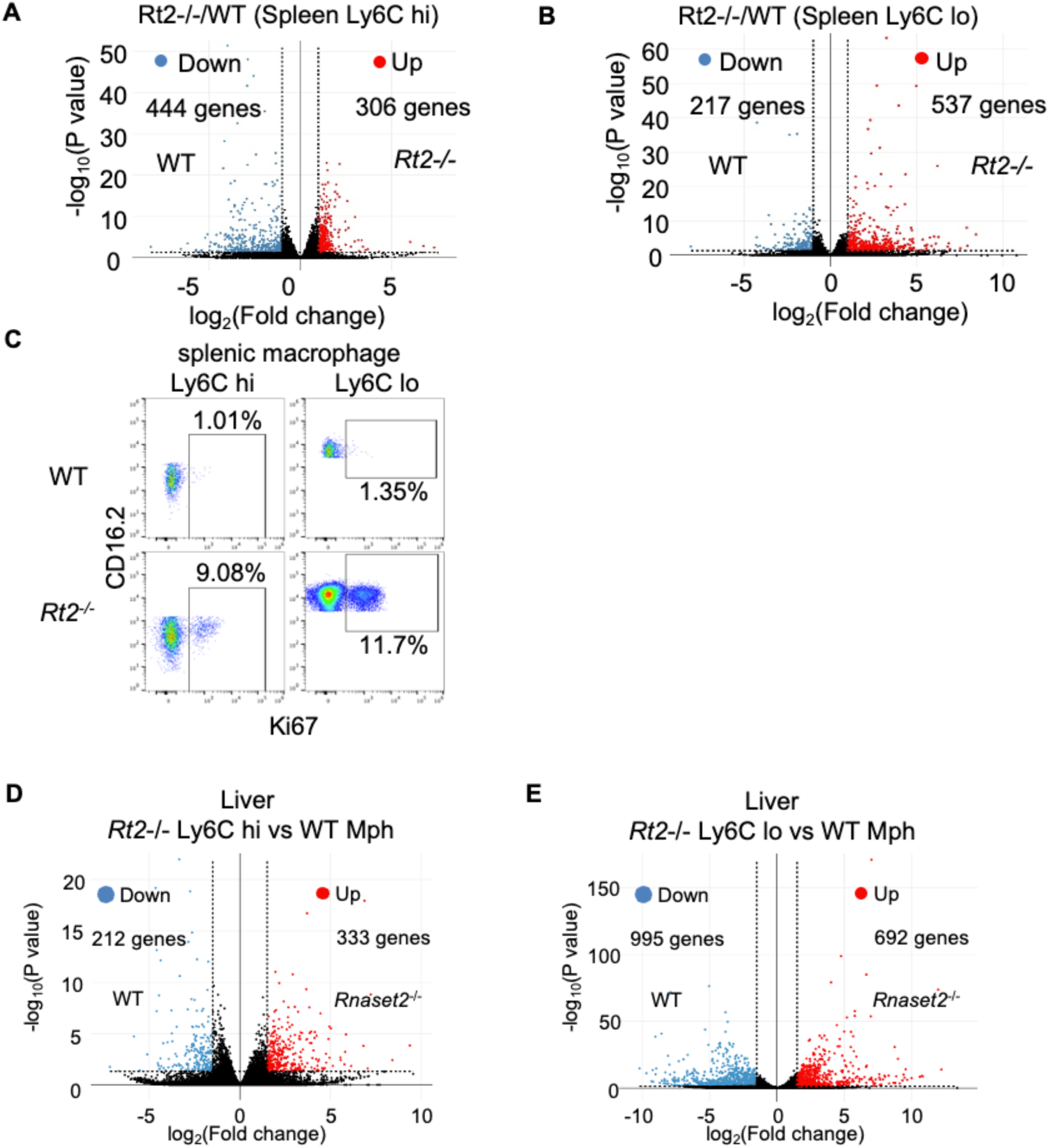
RNA-seq analyses of *Rnaset2^−/−^* mice. (A, B, D, and E) Volcano plots displaying log2 fold change of expression (X-axes) and log10 normalized expression (Y-axes) for the comparisons of splenic *Rnaset2*^−/−^ Ly6C^hi^ macrophages vs. WT Ly6C^hi^ macrophages (n = 3) (A); splenic *Rnaset2*^−/−^ Ly6C^lo^ macrophages vs. WT Ly6C^lo^ macrophages (n > 3) (B); hepatic *Rnaset2*^−/−^ Ly6C^hi^ macrophages vs. WT CD11b^+^ macrophages (n = 3) (D); and hepatic *Rnaset2*^−/−^ Ly6C^lo^ macrophages vs. WT CD11b^+^ macrophages (n = 3) (E). Genes with > 1.5-fold upregulated and downregulated expression are shown in red and blue, respectively. (C) Dot plots show the expression of Ki67 and CD16.2 in splenic macrophages from WT and *Rnaset2*^−/−^ mice.

**Fig. S4.**
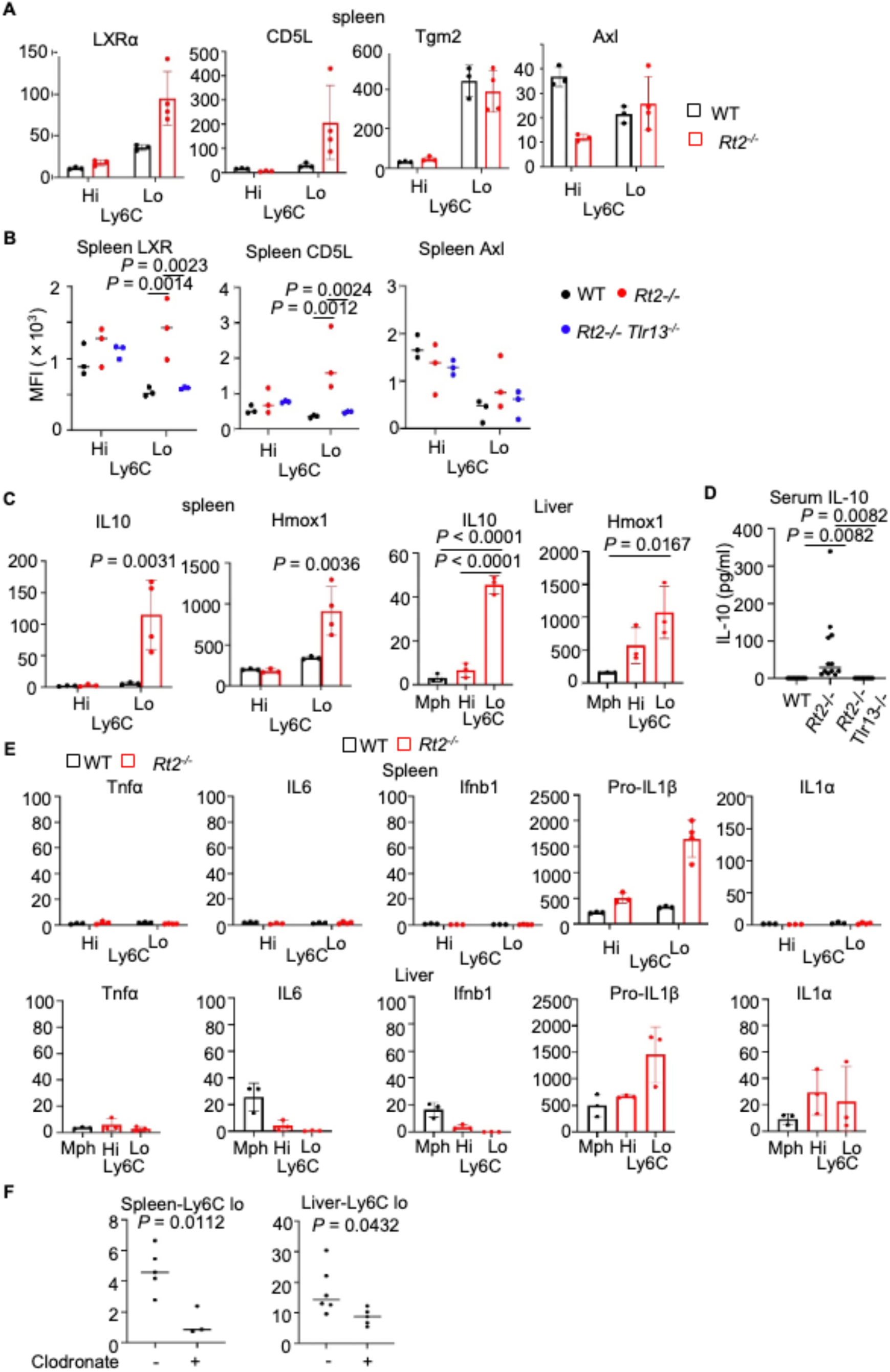
Gene expression in *Rnaset2*^−/−^ mice. (A, C, and E) Bars show reads per million (RPM) of indicated genes in indicated macrophage subsets in the spleen and liver. Mph indicates CD11b^+^ macrophages (n > 3). (B) Mean fluorescence intensity of indicated proteins in splenic Ly6C^hi^ and Ly6C^hi^ macrophages from indicated mice. (D) Serum levels of IL-10 in WT, *Rnaset2*^−/−^, and *Rnaset2*^−/−^ *Tlr13*^−/−^ mice. (n = 12). (F) Dot plots show the percentage of splenic and hepatic Ly6C^lo^ macrophages in WT mice treated with clodronate (25 mg/mice) at 16 h prior to analysis.

**Fig. S5.**
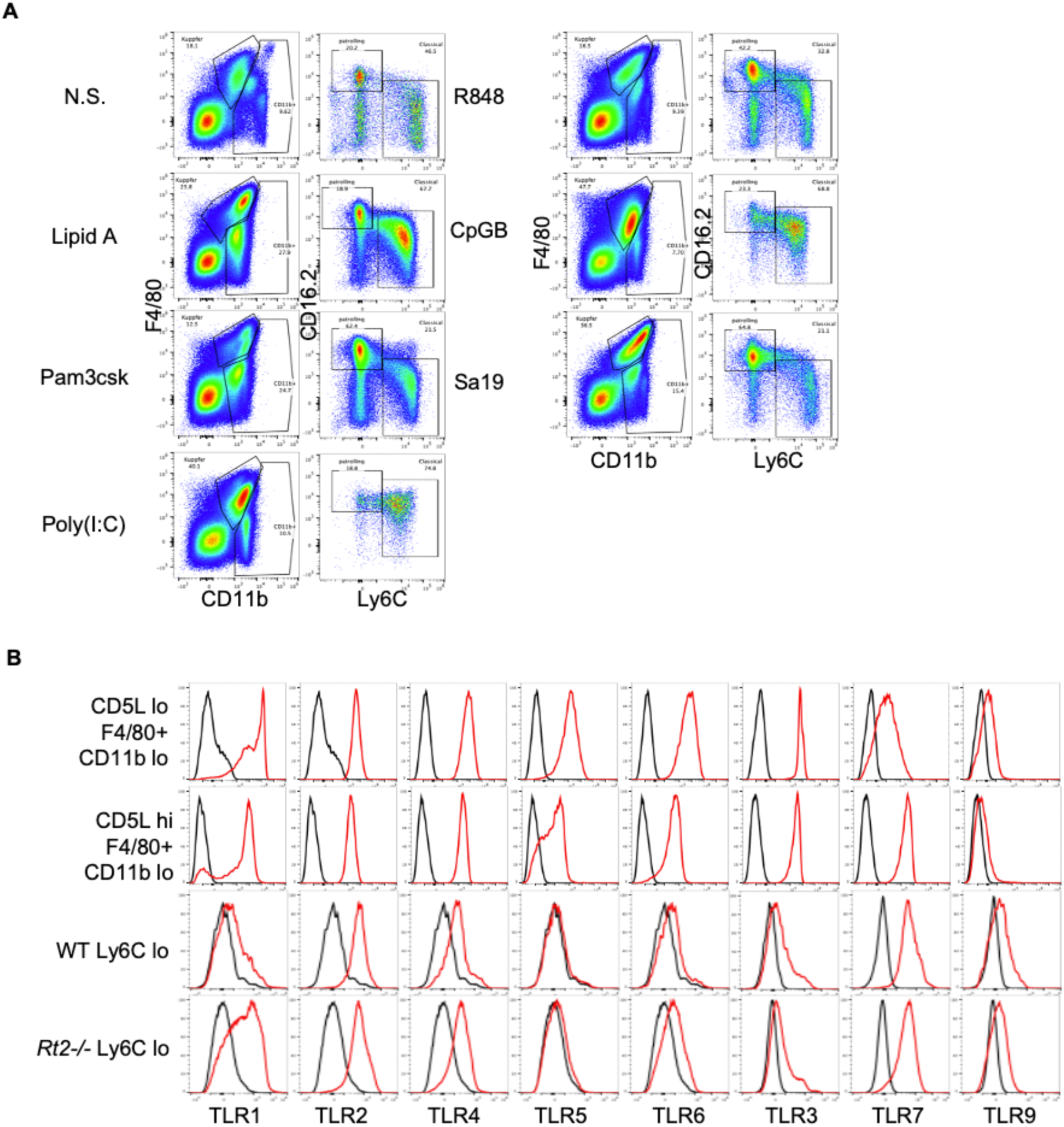
Effect of TLR ligands on hepatic macrophages. (A) Wild-type mice were administered TLR ligands, such as LPS (TLR4/MD-2), Pam3CSK4 (TLR1/TLR2), poly(I:C) (TLR3), R848 (TLR7), and CpG-B (TLR9), on days 1 and 2. Dot plots show hepatic macrophages in wild-type mice on day 3. (B) Histograms show staining of TLRs with antibodies against each TLR (red) and isotype-matched antibody (black).

**Fig. S6.**
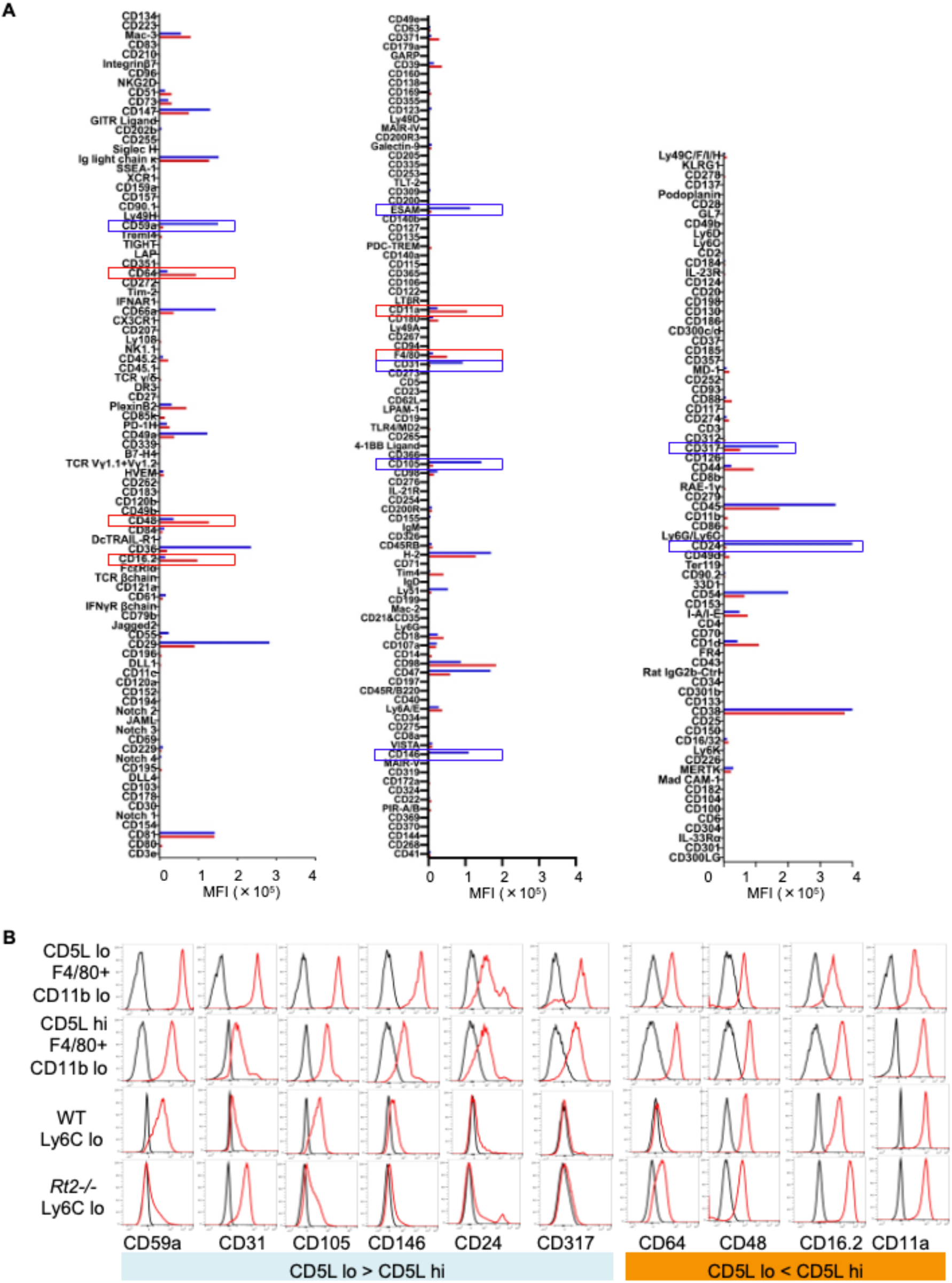
Cell surface marker expression in hepatic macrophage subsets. (A) Mean fluorescence intensity of indicated markers in KCs (blue) and KC2-like macrophages (red) from WT mice. (B) Red histograms show staining of indicated cell surface markers; black histograms show staining with isotype-matched antibody.

